# Computational modeling of left ventricular flow using PC-CMR-derived four-dimensional wall motion

**DOI:** 10.1101/2024.08.27.609991

**Authors:** Seyed Babak Peighambari, Tanmay Mukherjee, Emilio A. Mendiola, Amr Darwish, Lucas H. Timmins, Roderic I. Pettigrew, Dipan J. Shah, Reza Avazmohammadi

## Abstract

Intracardiac hemodynamics plays a crucial role in the onset and development of cardiac and valvular diseases. Simulations of blood flow in the left ventricle (LV) have provided valuable insight into assessing LV hemodynamics. While fully coupled fluid-solid modelings of the LV remain challenging due to the complex passive-active behavior of the LV wall myocardium, the integration of imaging-driven quantification of structural motion with computational fluid dynamics (CFD) modeling in the LV holds the promise of feasible and clinically translatable characterization of patient-specific LV hemodynamics. In this study, we propose to integrate two magnetic resonance imaging (MRI) modalities with the moving-boundary CFD method to characterize intracardiac LV hemodynamics. Our method uses the standard cine cardiac magnetic resonance (CMR) images to estimate four-dimensional myocardial motion, eliminating the need for involved myocardial material modeling to capture LV wall behavior. In conjunction with CMR, phase contrast-MRI (PC-MRI) was used to measure temporal blood inflow rates at the mitral orifice, serving as an additional boundary condition. Flow patterns, including velocity streamlines, vortex rings, and kinetic energy, were characterized and compared to the available data. Moreover, relationships between LV wall kinematic markers and flow characteristics were determined without myocardial material modeling and using a non-rigid image registration (NRIR) method. The fidelity of the simulation was quantitatively evaluated by validating the flow rate at the aortic outflow tract against respective PC-MRI measures. The proposed methodology offers a novel and feasible toolset that works with standard PC-CMR protocols to improve the clinical assessment of LV characteristics in prognostic studies and surgical planning.

## 1 Introduction

Intracardiac hemodynamics plays a crucial role in cardiac pumping efficiency, yet the intricate patterns of atrioventricular blood flow are not fully understood [1]. Functional hemodynamic markers such as stroke volume, ejection fraction, and mean arterial pressure, among others, provide important insight into the heart’s overall pumping function. These *organ-level* markers are often complemented by *tissue-level* measures of myocardial wall motion that can directly identify impaired wall contraction. However, the investigation of the potential presence of *regional* abnormal flow patterns within the left ventricle (LV) further complements *global* markers that may occur even prior to the manifestation of structural myocardial alterations [2, 3, 4]. This exploration advances the diagnostic capabilities, traditionally reliant upon functional assessments, towards early-stage identification and prediction of cardiovascular diseases (CVDs) [5]. Subsequently, there has been a growing significance placed on the individual-specific modeling and quantification of ventricular flow characteristics in assessing structural heart diseases and understanding flow-related mechanisms leading to CVDs [6, 7]. The characterization of vortical structures of the flow is a complex aspect of intracardiac fluid dynamics that has been indicated to be associated with cardiac pathologies [8]. The study of vortical structures, specifically vortices and vortex rings, in the LV during diastolic filling has been recognized as a potentially innovative method for analyzing the efficiency of LV pumping and the mechanosensitive remodeling of the heart [9, 10] and could impart a better understanding of the flow drivers of disease progression. A vortex is essentially a swirling flow of particles around a central axis, while vortex rings, a particular subcategory of vortices, are toroidal flow structures formed when fluid circulates around an imaginary ring-shaped closed surface [11, 12]. During diastole, the laminar blood flow from the left atrium to the LV is transformed into a vortex at the tips of the mitral valve leaflets. The formation of this vortex helps preserve the fluid momentum, enabling a smooth rerouting of the flow towards the aortic outflow tract with minimal turbulence generation [13]. This process helps to prevent significant losses of kinetic energy and increase LV ejection efficiency [14].

Recently, state-of-the-art methodologies have been introduced to evaluate blood flow dynamics in the cardiovascular system [15]. Several of these methods involve in-vivo techniques that directly measure the flow field through diagnostic tools such as Doppler imaging or particle image velocimetry [8, 16, 17], as well as conventional phase-contrast magnetic resonance imaging (PC-MRI) [1, 9, 18]. Despite their advances, these diagnostic techniques still offer a restricted view of cardiac functionality, with the diagnoses leveraging only a minimal portion of the data these techniques can provide [19]. Additionally, there are in-silico studies that integrate imaging measurements with computational modeling to analyze intracardiac fluid dynamics, employing two main modeling approaches. The first method computes the heart’s endocardium motion by modeling the entire electrophysiological structure of the myocardium, and from this basis, it predicts the flow within the ventricles [20, 21]. Despite the inherent benefits of constitutive-modeling-based approaches in recapitulating the various kinematic modes of cardiac contraction [22], they remain computationally cumbersome and may appear superfluous when the study focuses solely on flow behavior [2]. In contrast, the second method tracks the endocardial surface using time-resolved imaging modalities and applies this motion to fluid, resulting in a moving-wall CFD model [23, 24]. This approach requires considerably less in-vivo data, complexity, and computational power. Therefore, the moving boundary method is considered a more pragmatic alternative when attempting to model patient-specific intracardiac hemodynamics [5].

Various modalities and imaging methods have been used to capture the motion of the moving endocardium through a moving boundary patient-specific modeling approach. While there have been only a few studies using echocardiographic images to reconstruct the geometry (due to the low spatial resolution) [25, 26], cardiac computed tomography (CT) scans and cardiac magnetic resonance (CMR) images are the prevailing modalities used in patient-specific modeling. Cardiac CT scans provide excellent spatial resolution, which is beneficial in accurate reconstruction of the LV geometry. However, even with the recent advances in CT capabilities in the past few decades, the average temporal resolution of CT images cannot sufficiently estimate cardiac wall velocities for robust imaging-based assessment [27]. Moreover, high-resolution CT scans require high radiation dosage levels [28], increasing the risk imparted to patients with its clinical use. On the other hand, the order of CMR temporal resolution is significantly higher, which makes CMR a notably superior option for cardiac strain detection, endocardial motion tracking, and, consequently, a better choice for cardiac image-driven modeling [29]. The choice of imaging modality is crucial because CMR is a standard cardiac imaging modality and is commonly used in the clinical setting; however, one of the critical challenges with CMR-based moving boundary modeling is image registration and high-fidelity capture of geometry and wall motion, which is not as critical in dynamic CT-based modeling due to its inherently high spatial resolution. In addition, other modalities of CMR, such as PC-MRI, can be leveraged to personalize cardiac models further. PC-MRI allows for the non-invasive measurement of the complete set of blood flow velocity components in all four spatiotemporal dimensions of the heart [30]. While the clinical use of PC-MRI is well established [30], the integration of volumetric flow analysis from PC-MRI and moving boundary CFD methods has not yet been presented.

In this work, we implement a novel non-rigid image registration (NRIR) framework to evaluate the structural deformation of the myocardial wall from CMR images. The resulting endocardial wall displacements were imposed onto the individual-specific geometry integrated with the personalized inflow boundary condition assessed at the mitral valve orifice using PC-MRI modality to numerically model the intraventricular flow through moving boundary CFD simulations. Our method extends upon previous CMR-based CFD modeling, introducing an innovative non-rigid estimation of 4D myocardial deformation, eliminating the need for complex mathematical modeling while precisely tracking the wall. In addition, PC-MRI was used for the first time in LV hemodynamic modeling, in conjunction with standard cine CMR, to impose the spatiotemporal blood flow rate at the mitral inflow track, eliminating the use of other modalities to capture the flow. In addition to characterizing intracardiac LV hemodynamics, PC-MRI data at the aortic outflow track was used to evaluate the ejection flow rate and distribution predicted by the method. The intraventricular 3D velocity streamlines, vortex rings, planar vorticity, and kinetic energy were analyzed in a cardiac cycle. This approach provides a platform to further investigate the correlation between the wall kinematics and flow characteristics.

## 2 Materials and methods

### 2.1 CMR image acquisition

Cine- and PC-MRI imaging were performed for a patient presenting mitral valve prolapse (MVP) (n = 1). The standard functional assessment using cine imaging involved the acquisition of nine short-axis (SA) image planes between the atrioventricular junction and the apex of the LV and a single four-chamber plane using a steady-state free-precession sequence. Additionally, PC-MRI images were acquired in the SA planes of the aortic and mitral valve planes. CMR and PC-MRI images were obtained using the same protocol in a 3.0T clinical scanner (Siemens Verio; Siemens, Erlangen, Germany) with phased-array coil systems. The scan parameters were as follows: flip angle of 65° - 85°; repetition time of 3.0 ms; echo time of 1.3 ms; in-plane spatial resolution of 1.7-2.0 × 1.4-1.6 mm; slice thickness of 6 mm with 4 mm interslice gap; and temporal resolution of 35-38 ms, which lead to 20 timeframes per cardiac cycle. Following image acquisition, all personal identifiers were removed per the National Institute of Health de-identification protocol, comprising 18 elements considered protected health information [31]. The research protocol was approved by the Houston Methodist Research Institute review board.

### 2.2 CMR-derived motion analysis

Cardiac motion was quantified for the duration of a single cycle using a stack of the nine SA images of the LV. Segmentation was performed to isolate the endo- and epicardium of the LV using two-dimensional (2D) contours that were semi-automatically drawn on each image plane using Segment [32]. The pixels in the space between the two 2D contours in each image were resampled along the through-plane axis using Delaunay triangulation and smoothed through the application of radial basis functions, thus providing a three-dimensional representation of the LV anatomy. The segmentation was repeated for each time frame in the sequence of twenty images describing the cardiac motion, and the resulting images were subjected to NRIR using the diffeomorphic demons algorithm. Our framework [33, 34, 35] extends upon the classical optimization-based registration problem to find the optimal transformation in aligning a reference frame, ℱ, and a moving image, *M* through the minimization of the cost function, *W* as

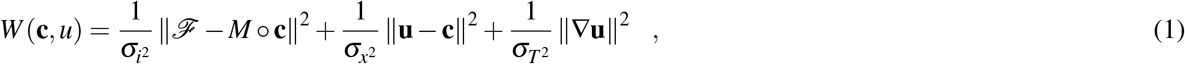

where *σ*_i_ and *σ*_x_ are the noise intensity parameter and spatial uncertainty parameter, respectively, and *σ*_T_ is a regularization factor. Here, the parametric transformation (**u**) refers to the Cartesian displacement vector at each pixel between two consecutive time frames, and **c** denotes the corresponding non-parametric spatial transformation.

#### Myocardial strain estimation

The cardiac motion was additionally quantified by estimating myocardial strains from the displacements derived through NRIR. The myocardial strains were calculated at end-systole (ES) with respect to end-diastole (ED), thus describing the contraction phase of the cardiac cycle. We used large deformation theory to derive the Green-Lagrange strain tensor (**E**), processing the Cartesian displacements obtained from NRIR as pixel-wise data. The strain computation involved the calculation of the total deformation gradient (**F**) as the cumulative or multiplicative deformation gradient across all time frames such that,

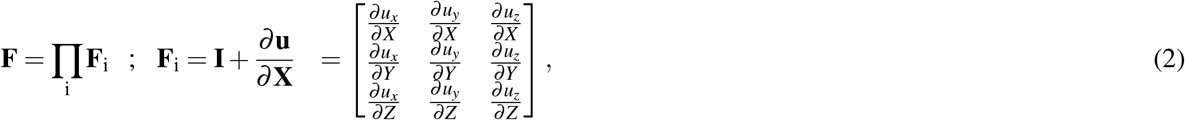

where **F**_**i**_ describes individual deformation gradients obtained using the displacements between two successive time frames, *x, y, z* and *X,Y, Z* represent the Cartesian axes in the moving and fixed frames, respectively. In this context, u_x_, u_y_, and u_z_ denote the displacements in the x, y, and z directions, respectively, between successive time frames. **F** was used in conjunction with the identity matrix (**I**) to express **E** as: 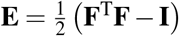.

### 2.3 Implementation of the moving boundary CFD model

The segmented myocardial geometry was used to isolate the contours describing the endocardium in each SA image at ED. The points corresponding to the endocardium in Cartesian space were triangulated using the Delaunay triangulation algorithm, thereby providing a surface geometry of the endocardium (Fig. 1A). This surface formed the initial state for the subsequent CFD analysis, and the displacements obtained through NRIR from the succeeding time points were mapped onto it using a closest neighbor least-squares approximation. Additionally, the displacements were trilinearly interpolated to increase the number of time frames from 20 to 321. The fluid volume within the endocardium was discretized using linear tetrahedral elements and dynamic mesh methods at ED. A laminar viscous model was implemented for the fluid simulation. Blood was assumed to be incompressible, homogeneous, and, due to high shear rates in the LV, Newtonian. The blood density and viscosity were set to 1040 *kg*/*m*^3^ and 0.0035 *Pa · s*, respectively [36]. A no-slip condition was specified between the fluid and the LV wall. The coupled pressure-volume solver was implemented to accommodate the large displacements of the LV endocardium [37]. The endocardial displacements acquired from the NRIR method were imposed on the discretized moving wall using a user-defined function (UDF) in Ansys Fluent 2020 R2. The UDF matched the displacement of the points on the LV segment to the closest corresponding node on the LV mesh. For the governing equation, the incompressible Navier–Stokes equations with the arbitrary Lagrangian-Eulerian (ALE) formulation were used [36], and the continuity and the momentum equations were defined as:

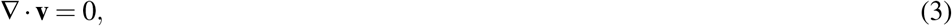

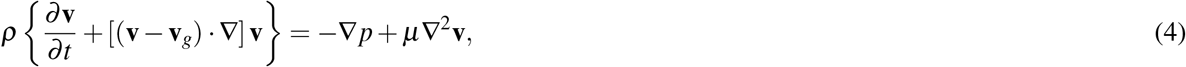

where **v** is the flow velocity,**v**_*g*_ is the cordinate velocity, *p* is the total pressure, and *ρ* and *µ* are the blood density and viscosity, respectively.

**Fig. 1.**
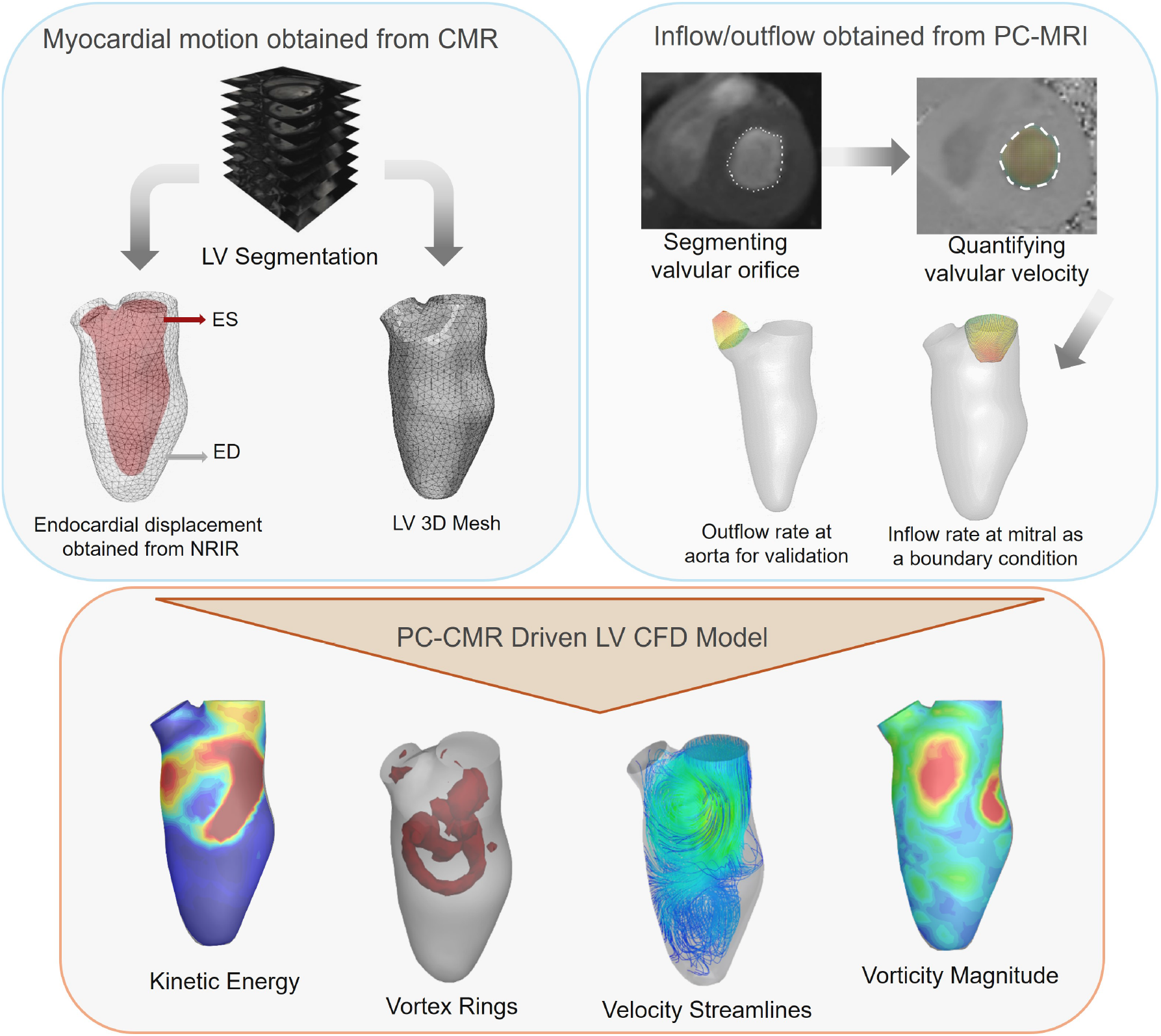
Schematic of the proposed cardiac magnetic resonance PC-MRI imaging-based Computational fluid dynamics (CFD) setup. Section (a) illustrates the reconstruction of LV geometry using short-axis(SA) CMR images and the non-rigid image registration (NRIR) method to obtain the endocardial wall displacement from end-diastolic and end-systolic volumes. Section (b) depicts the flow rate acquisition by integration of PC-MRI flow velocity measurements over the mitral and aortic orifices

### 2.4 PC-MRI derived flow analysis

PC-MRI is a non-invasive imaging method capable of assessing the 4D velocity of blood flow within the heart and major blood vessels and has been in routine clinical use for several decades [38, 39]. While conventional CMR primarily represents the regional signal amplitude at each pixel in the image, PC-MRI utilizes the phase shifts induced by moving protons (spins) of the MR signal to detect motion and velocity. In clinical practice, it is expected to map the velocity component of PC-MRI perpendicular to a 2D plane for volume flow measurements [30, 40]. In this study, the blood flow velocity field was calculated at the 2D area of the mitral inflow tract in 12 time points during diastole to tailor our approach to the same protocol used in the clinical setting (Fig. 2a). At early diastole and with the onset of inflow E-wave, the mitral valve was fully open, and the shape of the area of the inflow velocity field was nearly circular at time points T_1_ to T_3_, while for the rest of the diastole, as the velocity dropped, the area of the inflow velocity field became amorphous (Fig. 2a). Then, the mitral velocity field was integrated over the changing mitral orifice to acquire the flow rate; the flow rate measured for the cardiac cycle was applied as a transient flow boundary condition in the CFD model (Fig. 2b). The early mitral inflow rate peaked between time points T_2_ and T_3_ at −322 mL/s, and the late inflow rate (occurring during atrial filling) peaked at 147 mL/s at time point T_11_ (Fig. 2b). Volumetric flow at the aortic outflow tract was also captured and used to validate the CFD modeling approach; aortic flow is presented in Section 3.4.

**Fig. 2.**
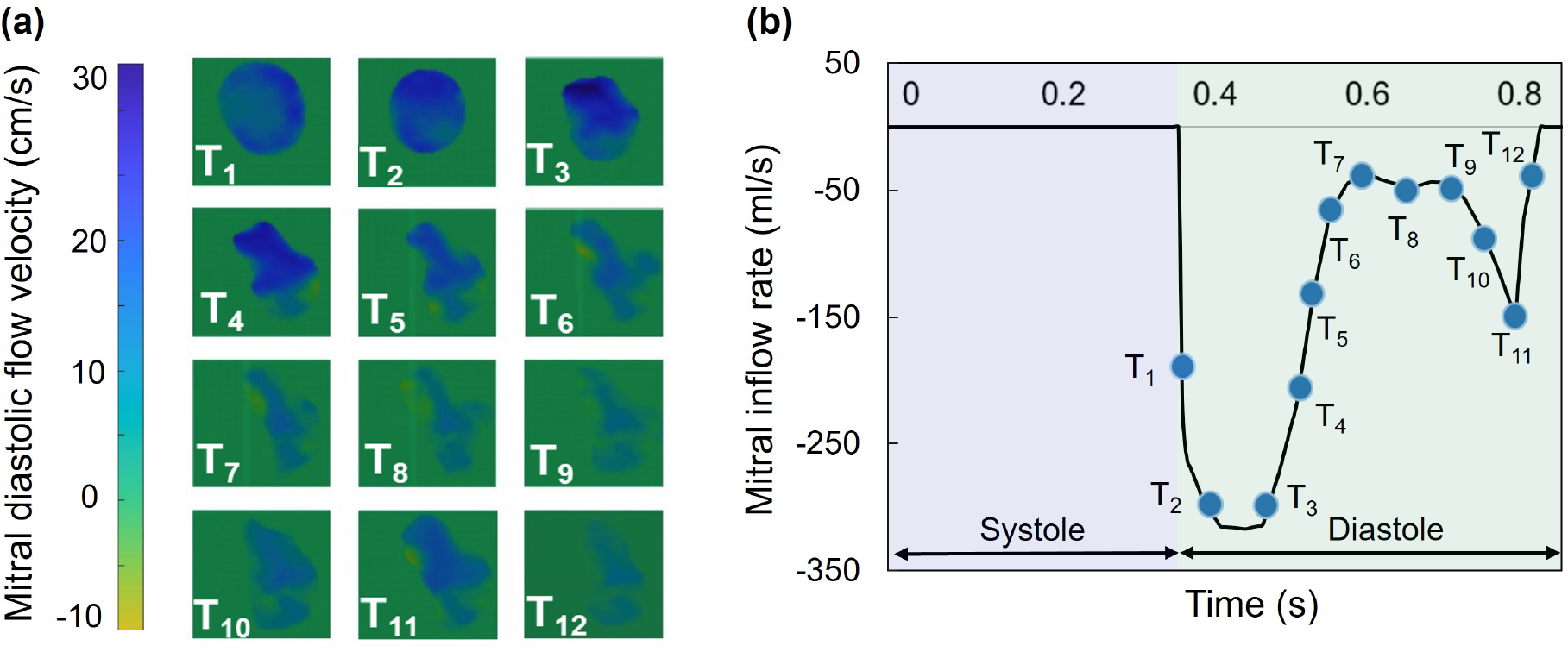
(a) 2D PC-MRI derived flow velocity at the mitral inflow track at 12 time points during the diastolic phase; T_1_ to T_4_ are in rapid filling, T_5_ to T_9_ are in diastasis, and T_12_ to T_10_ are in atrial filling(b) The mitral volumetric flow rate imposed on the CFD model in a cardiac cycle, derived by integrating flow velocity over the altering velocity area.

### 2.5 Hemodynamic characteristics quantification

Following the CFD simulation, several metrics were computed to characterize spatiotemporal LV hemodynamics: streamlines, vorticity magnitude, iso-surface vortex rings, and kinetic energy. There are multiple methods to identify vortical structures in cardiac hemodynamics; in this work, we chose to use the vorticity magnitude Ω, which is defined as:

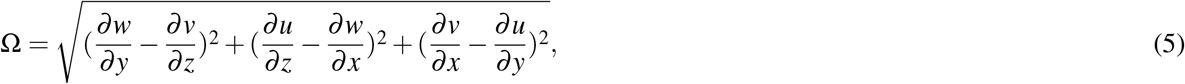

where *u, v*, and *w* represent blood flow velocity in the *x, y*, and *z* directions, respectively [41].

We additionally characterized vortices using the *Q* criterion method. This method visualizes vortex ring iso-surfaces by identifying vortices in regions where the second invariant of the velocity gradient (∇**v**), termed *Q*, is positive [42, 43]. The *Q* criterion is defined as:

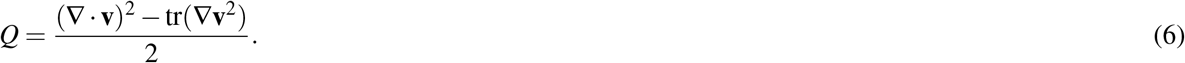

## 3 Results

### 3.1 Full cycle high-fidelity velocity streamlines and vorticity measurements

Velocity streamlines (Fig. 3) and vorticity (Fig. 4) were calculated throughout the entire cardiac cycle. Whereas the aortic outflow and systolic vorticity are driven by the endocardial wall motion captured through the methods presented in section 2.2, the flow at the mitral valve was imposed through PC-CMR at the atrial filling phase. In the diastolic phase, the highest inflow velocity reached 55 cm/s during rapid filling, while the inflow velocity reached 30 cm/s during atrial filling (Fig. 3). The highest velocity flow remained in the basal area of the LV between the outflow and inflow tracks throughout the entire diastolic phase (Fig. 3). Systolic flow velocity peaked at 85 cm/s at the atrial outflow track. As rapid diastolic filling transitions into diastasis, vorticity peaks (Fig. 4) with a volumetric averaged magnitude of 15.7 *s*^−1^ (Fig. S1), and focal concentration in the central basal area of the LV. In the systolic phase, prominent vortices were evident near the aortic outflow tract at the peak and late systole with a volumetric averaged peak magnitude of 12.5 *s*^−1^ (Fig. S1) focally concentrated surrounding the aortic valve annulus (Fig. 4).

**Fig. 3.**
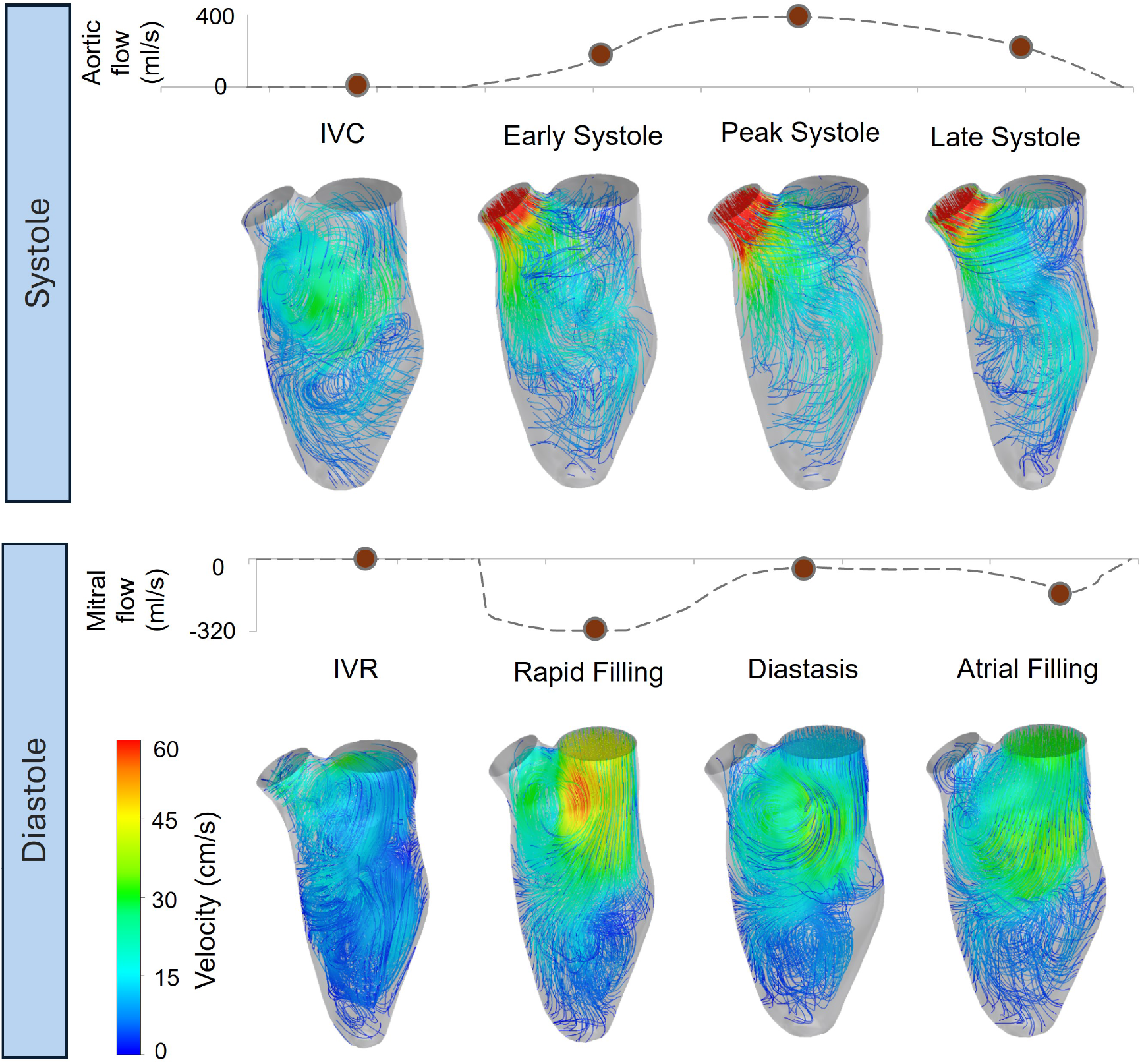
Top row: 3D velocity streamlines in the systolic phase at four time points: isovolumetric contraction (IVC), early systole, peak systole, and late systole. Bottom row: Velocity streamlines in the diastolic phase at four time points; isovolumetric relaxation (IVR), rapid filling, diastasis, and atrial filling. The E-wave jet, A-wave jet, and the counter-rotating vortices are captured.

**Fig. 4.**
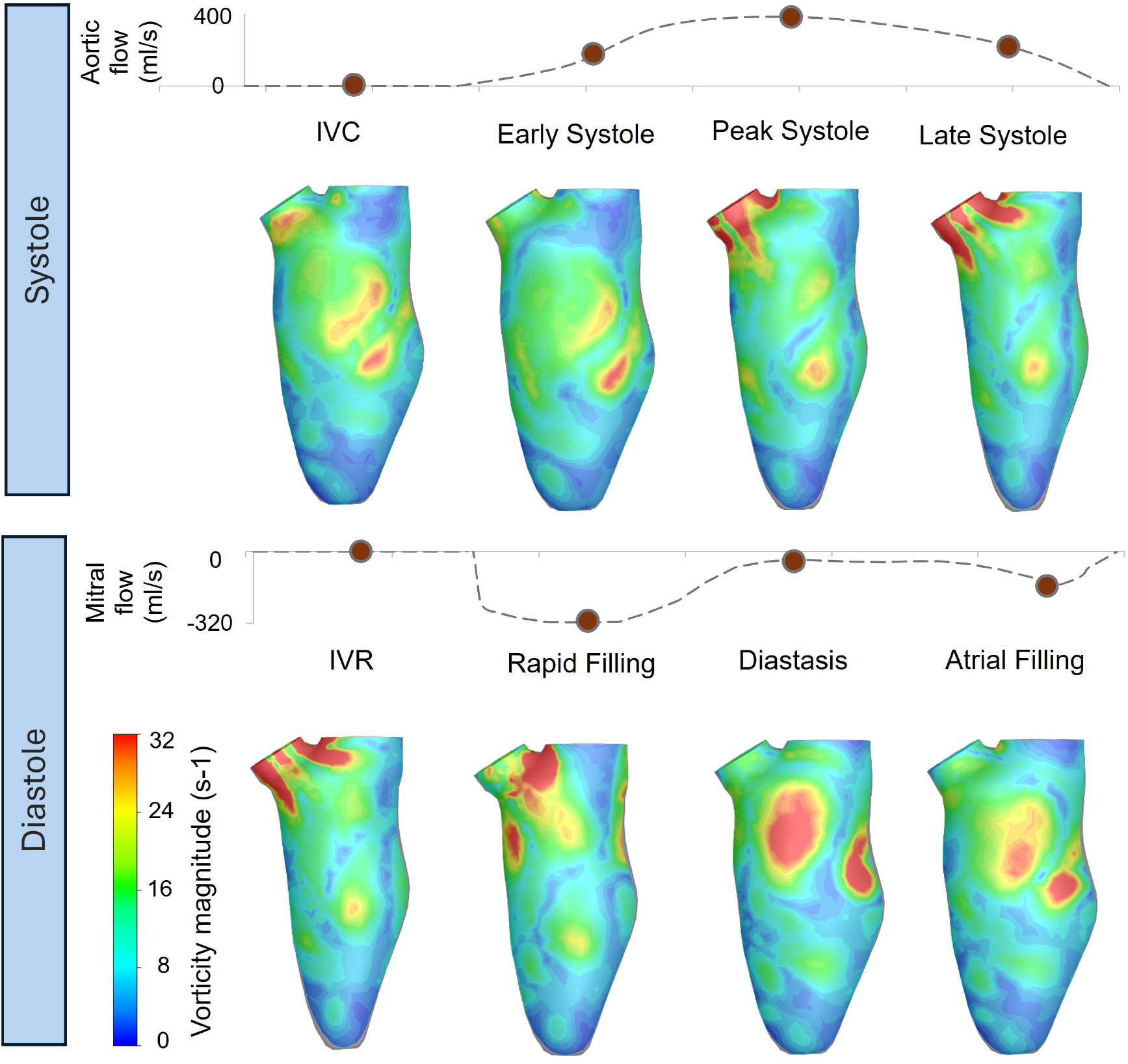
Vorticity magnitude presented in the coronal plane, crossing the mitral and aortic orifices. Top row: Vorticity magnitude in the systolic phase at four time points: isovolumetric contraction (IVC), early systole, peak systole, and late systole. Bottom row: Vorticity magnitude in the diastolic phases at four time points; isovolumetric relaxation (IVR), rapid filling, diastasis, and atrial filling.

### 3.2 Characterization of intraventricular vortex rings at high spatiotemporal resolution

The LV vortex rings were captured in the cardiac cycle (Fig. 5). We found setting *Q* = 50 (Eqn. 6) provided the optimal vortex visualization. Two main eddy currents were observed forming in the diastolic phase and shrinking in the systole phase. At the onset of the first diastolic jet flow (E-wave; rapid filling), the biggest vortex ring forms downstream of the mitral inflow tract, and as diastasis progresses, this vortex flow gradually shifts towards the midsection of the LV. During the second peak of the diastolic phase (A-Wave; atrial filling), a secondary vortex emerges near the inlet valve and extends downstream. The vortices reached their maximum intensity at the end of the diastole. Aggregation of the vortex ring in the mid-section can likely be attributed to the interactions between the impinging jet of flow and the relatively stagnant septal flow. At early systole, the outflow jet forms a vortex ring encircling the aortic outflow tract, while during the rest of the systolic phase, the LV vortex rings structured in the diastole are pushed out with the outflow.

**Fig. 5.**
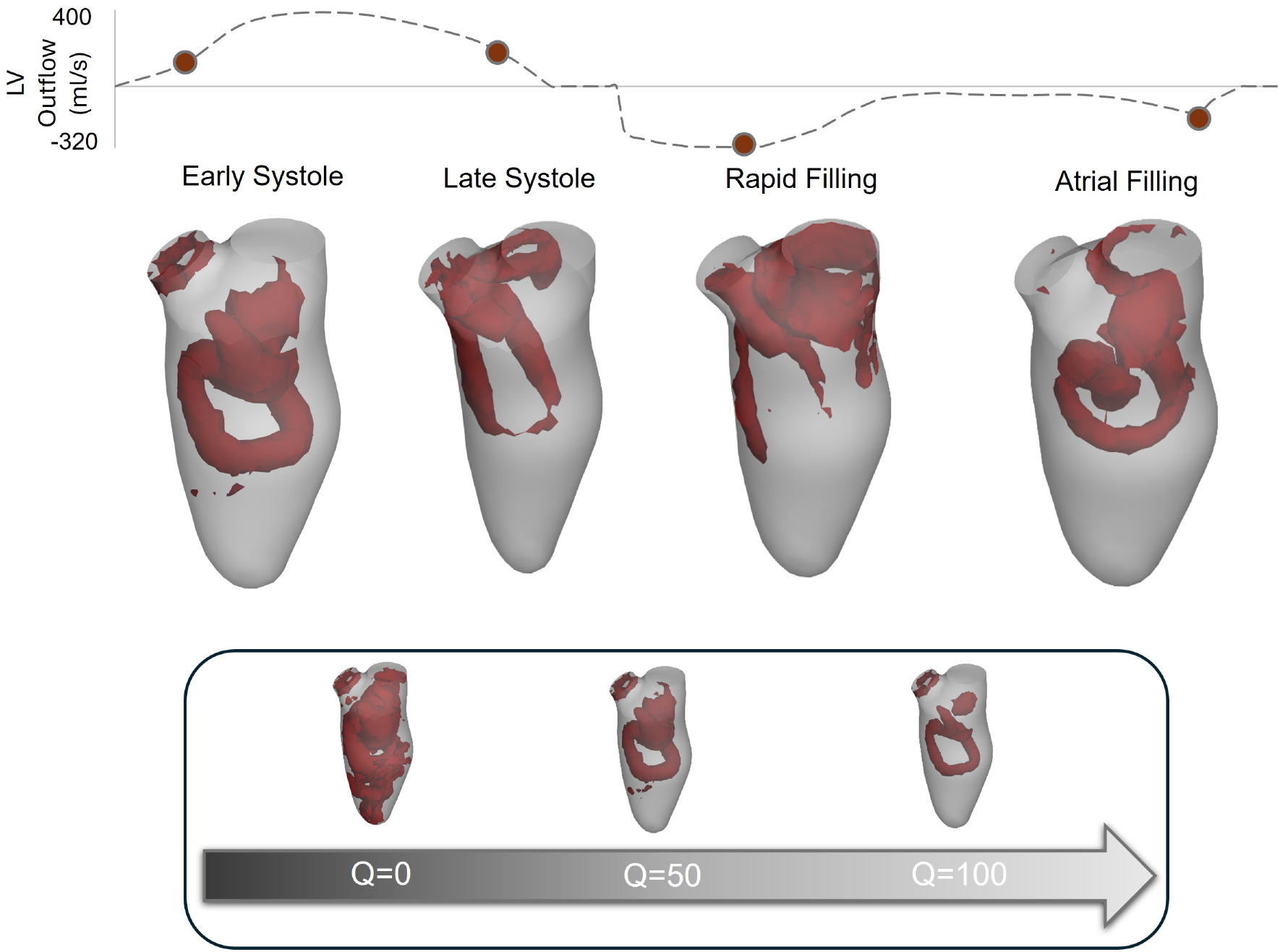
Isosurface of vortex rings illustrated with *Q* criterion method, where *Q*=50. Three main vortex rings correspond to the three jets of flow in the cardiac cycle. The first was created at the early systolic phase, encircling the aortic outflow jet. Second, the vortex ring developed with the early filling jet of inflow, which was the largest. Third, the vortex ring was built with the onset of atrial filling.

### 3.3 Relationship of intraventricular flow kinetic energy and myocardial strains

Kinetic energy was averaged in the entire LV cavity and also examined at three SA cross-sections at the basal, mid, and apical levels of the LV (Fig. 6a). The average kinetic energy of each of the three regions, plus the average over the entire LV volume, was calculated throughout the entire cardiac cycle (Fig. 6b). The increased kinetic energy in the LV cavity during the systolic phase was dominated by behavior at the basal plane. Kinetic energy reached a minimum at IVC and quickly peaked at the early diastole. At all time points throughout the cardiac cycle, the kinetic energy of the fluid was less at each SA level, moving from the base to the apex. Moreover, after obtaining the plane-averaged vorticities in the same SA planes (Fig. S2), we analyzed the correlation between the kinetic energy and vorticity during the cardiac cycle at nine evenly spaced time intervals (Fig. 6c). In addition, the relations between the kinetic energy and the longitudinal, radial, and circumferential strains of the myocardium wall acquired from the NRIR framework were examined. Mean strains and kinetic energy are presented on the basal SA plane (Fig. 7a) at eight evenly spaced time intervals during the systolic phase. Radial and circumferential strain were positively correlated with kinetic energy (Fig. 7e,f), whereas longitudinal strain did not show a significant correlation (Fig. 7d).

**Fig. 6.**
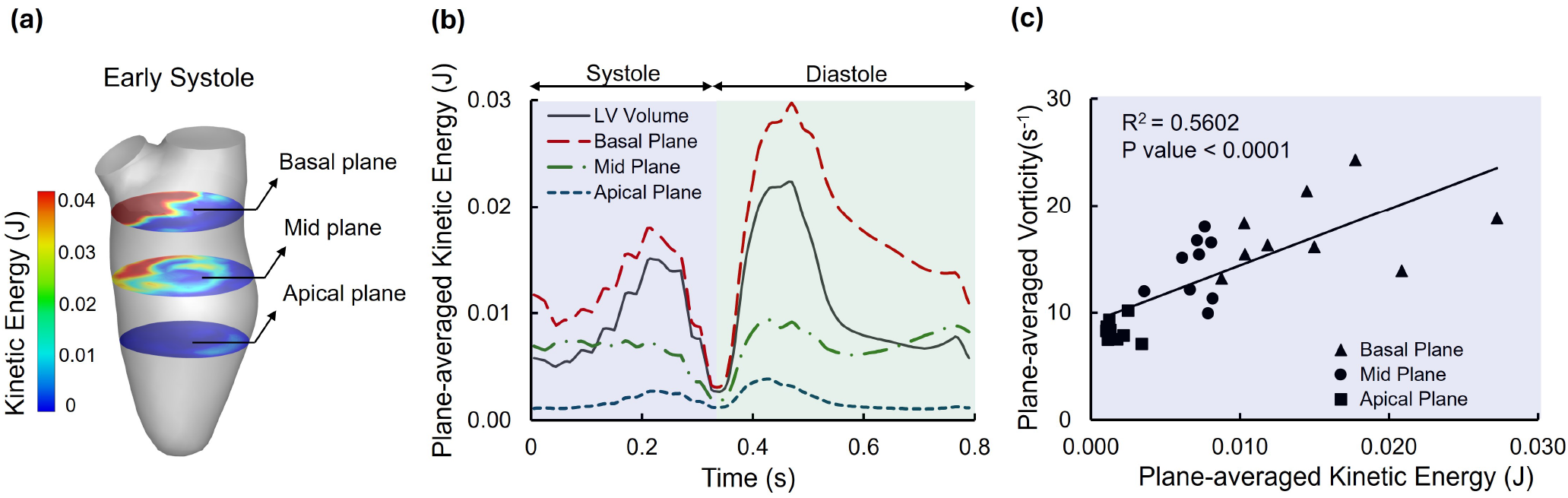
(a) Kinetic Energy field at early systole phase in three short-axis planes of basal, mid, and apical sections. (b) Kinetic energy averaged in the LV volume and the three short-axis planes during a cardiac cycle. (c) Vorticity magnitude averaged in the LV volume and three short-axis planes during a cardiac cycle. (d) the correlation between the kinetic energy and vorticity magnitude in each short-axis plane during eight evenly spaced time intervals.

**Fig. 7.**
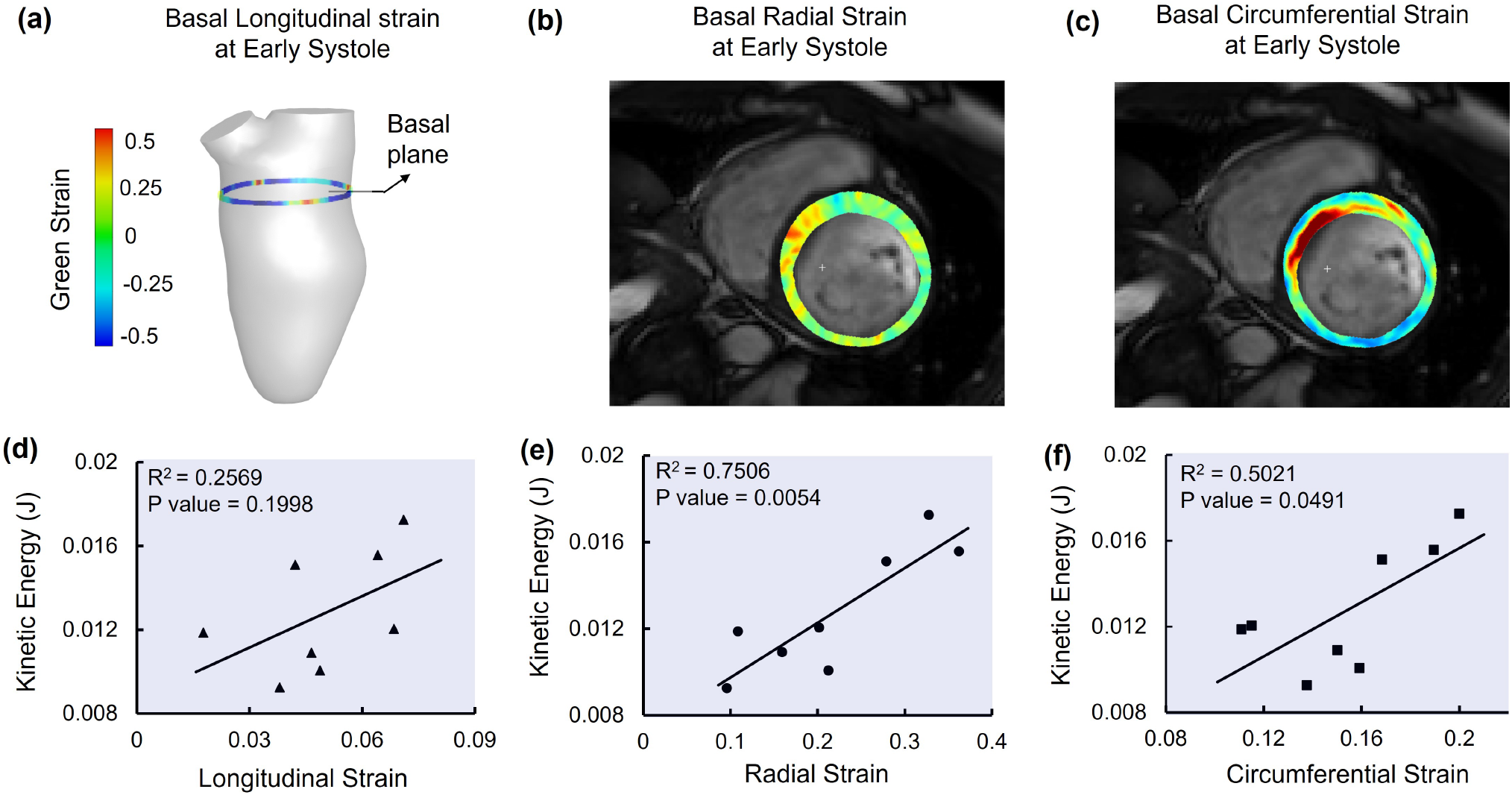
(a) Longitudinal strain demonstrated at the base of LV simulated model at early systole (b) Early systolic radial strain at the base of LV demonstrated in CMR short axis view (c) Early systolic circumferential strain at the base of LV demonstrated in CMR short axis view. (d-f) The correlation of systolic longitudinal, radial, and circumferential strain with kinetic energy at the basal level, respectively.

### 3.4 Validation against PC-CMR aortic flow measurements

To validate the simulated results, the flow velocity and the volumetric flow rate crossing the aortic outflow tract obtained from the simulation was compared to the PC-MRI results (Fig. 8a). As inlet flow rate and wall motion were prescribed as inputs to the CFD model, the outflow behavior was ideally suited to serve as a measure of the accuracy of the overall behavior of the model. The LV CFD model was able to closely estimate the PC-MRI-derived aortic outflow rate throughout the entire cardiac cycle (Fig. 8b). The root mean square error (RMSE) over one systolic phase was 37 mL/s, and the maximum error, which occurred near peak systole (T3 time point), was 52 mL/s. It should be noted that the volume flow rate derived from PC-MRI was captured at the aortic outflow tract downstream of the aortic valve. Therefore, there is a continued flow after the end of the systolic phase in the aorta, which was not considered in the ventricular model.

**Fig. 8.**
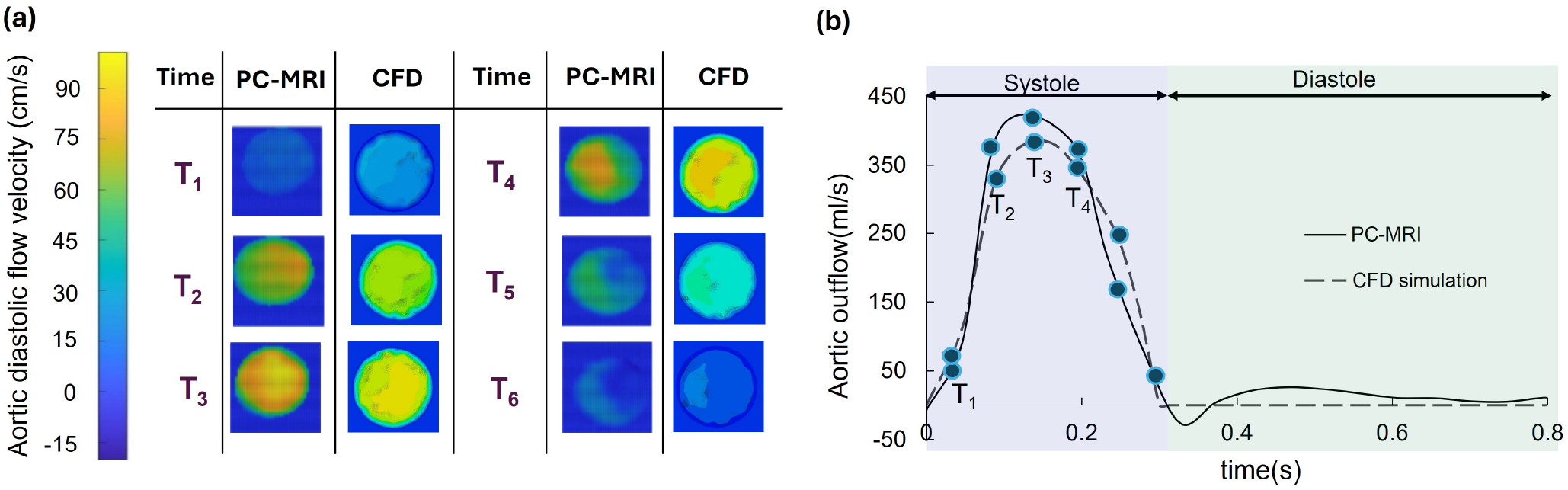
(a) 2D PC-MRI derived flow velocity compared to the CFD simulated velocity at the short axis view of the aorta at six time points during the systolic phase. (b) The aortic volumetric outflow rate assessed from PC-CMR compared to the aortic volumetric outflow rate obtained from the CFD model in one cardiac cycle

## 4 Discussion

### 4.1 Leveraging the combination of PC- and CMR to enhance LV flow characterization

Accurate patient-specific characterization of intracardiac hemodynamics is essential to improve understanding of the potential adaptive and maladaptive effects of altered flow in the LV. While PC-CMR provides comprehensive flow velocity data, it is challenged by cycle-to-cycle reproducibility in flow patterns and cannot resolve small-scale flow features [44]. However, incorporating in vivo flow data from PC-CMR with robust numerical modeling methods provides a method capable of characterizing high-resolution small-scale flow behavior. In this study, we presented a pragmatic, streamlined approach to patient-specific LV hemodynamic modeling by reducing the reliance on multiple clinical tests and decreasing computational cost. While previously devised methods of patient-specific hemodynamic modeling have been conducted with imaging-derived wall motion [5], no previous work has incorporated in vivo velocity data for initial boundary conditions and model validation. Whereas previous approaches make use of wall motion to derive inflow boundary conditions, the presented approach more closely models physiological conditions and provides for model verification through outflow comparison. Considering the increasing utilization of PC-CMR for volumetric flow measurements in clinical settings, as well as the higher temporal resolution in comparison to CT scans [29], the presented approach holds promise to enhance the fidelity of LV hemodynamic characterization without the computational cost of traditional CFD approaches.

### 4.2 Enabling high-fidelity quantification of intraventricular vortex development

After the CFD modeling, the LV vortical structures were examined using 3D velocity streamlines (Fig. 3), 2D vorticity magnitudes in the longitudinal plane (Fig. 4), and 3D vortex rings (Fig. 5). During systole, the constricting LV wall generated a jet of flow at the left ventricle outflow tract (LVOT) with a well-defined vortex ring surrounding the aortic annulus. LV flow studies have mainly focused on the characteristics of flow in the diastolic filling, and systolic vortices have been explored separately in the aortic arch. To our knowledge, there has been no previous report of a vortex ring surrounding the LVOT at early systole. During diastole, with the onset of rapid filling, the interaction between the E-wave inflow jet and the relatively stagnant IVR flow created a vortex ring encapsulating the mitral orifice (Fig. 5). At diastasis, the vortex ring penetrated further towards the LV apex and developed into two major counter-rotating vortices in the longitudinal view (Figs. 3 and 4). The A-wave inflow jet of atrial filling created a fragmented small vortex ring around the mitral orifice and extended the previous vortices to their maximum size by pushing them deeper into the LV. The diastolic vortex rings and magnitudes illustrated are consistent with existing literature [11, 45], demonstrating the robustness of the presented modeling methods. While an efficient diastolic flow in healthy cases could necessitate that a single vortex encloses the whole LV to redirect the flow from the mitral valve to the aortic valve and minimizes the loss of kinetic energy [6, 8], at least two significant vortices exist in our model. Multiple vortical structures in the LV have been reported before in patients with mitral valve disorders [46]. Although we had not modeled the mitral valve, the altered flow in the presence of MVP was detected, likely as a result of the incorporation of patient-specific LV structure and wall motion. Future studies using an extended version of the presented approach, including valves, could indeed separate the contribution of leaflet and wall motion to the flow behavior.

### 4.3 The interplay between LV flow dynamics and endocardial wall kinematics

Despite the potential for flow dynamics to drive mechanosensitive remodeling events, studies are yet to associate kinematic markers of cardiac contractility (strains) with functional markers of atrioventricular flow. A few studies have explored alterations of the kinetic energy of LV flow in diastole [1]; here, we have proposed the correlation of kinetic energy and vorticity during a complete cardiac cycle, gathering data from three layers of LV. Although flow streamlines penetrate the apex level, the observed vortices are centered at the base and mid-level. It can be inferred that the kinetic energy of the flow is being conveyed with the vortices and not with flow streamlines; our correlation analysis affirms this notion by displaying a significant correlation between kinetic energy and vorticity. While the correlation of kinetic energy with the E- and A-waves (i.e., diastolic flow) has been evidenced [9], the corresponding characterization of kinetic energy in systolic flow remains understudied. The presented method enables the exploration of the impact of altered myocardial wall motion without the cumbersome computation of coupled electromechanical models that capture the myocardial strains. By integrating the presented CFD method with the NRIR framework on PC-MRI images, it was possible to investigate the relation between myocardial strains and systolic kinetic energy. In addition to built-up diastolic momentum [47], a positive correlation between myocardial strains and displacements during systole was observed (Fig. 7). Interestingly, longitudinal strain, an established clinical kinematic marker of overall LV function, did not significantly correlate with systolic kinetic energy. This suggests a possible transition from longitudinally driven flow in health to flow that is regulated primarily by circumferential cardiac motion in disease. Indeed, subclinical shifts in fiber arrangement, which also contribute to heterogeneous contractile patterns [48, 49], may provide variations in the resulting systolic flow patterns. The presented modeling platform holds promise to elucidate such alterations in the relationship between wall motion and intracardiac flow behavior, leading to a better understanding of the determinants of disturbed flow in the diseased heart.

### 4.4 Limitations and future directions

In this study, the mitral and aortic valves were considered to be fully closed during systole and diastole, respectively; therefore, while the valves were closed, the regurgitant flow detected in the PC-CMR data was not considered in the CFD model (Fig. 8b). The inclusion of structures representing the mitral and aortic valves in the model could increase its robustness. Such an addition could allow for the study of the independent effects of valve and wall motion on ventricular flow. Also, as expected, the calculation methodology presented herein was influenced by the image quality, with all the images subjected to Gaussian smoothing to regularize displacements. Image gradients along the tissue-pool interface of the endocardium are in constant flux due to the rapid movement of the myocardial wall, with these gradients generating unnaturally high pixel transformations (displacements). New imaging methodologies are needed to improve the spatial resolution of CMR scans, reduce the dependence on image filtering, and minimize overall anomalies.

## 5 Conclusions

There remains a pressing need for a more robust image-based approach to study the role of LV hemodynamic simulations in the prediction and detection of cardiovascular diseases (CVDs). This approach should aim to reduce the computational burden of simulations while maintaining the accuracy required to capture critical clinical hemodynamic characteristics. The proposed method in this study reinforces the concept that when standard (PC)-CMR imaging is combined with CFD modeling, it can provide a cost-effective and dependable method for obtaining high-quality results in cardiac research. This combination not only enhances the temporal accuracy of solid aspects of the LV (endocardial movement) but also provides additional insights into fluid dynamics (PC-CMR), which leads to a more practical solution for advancing our understanding of cardiovascular diseases.

## 6 Acknowledgements

R.A. was supported by the NHLBI grant R00HL138288. T.M. was supported by the American Heart Association (AHA) predoctoral fellowship 24PRE1240097.

## 8 Data availability

The data that support the findings of this study are available from the corresponding author, R.A., upon request.

## 9 Supplementary material

**Supplementary Figure S1.**
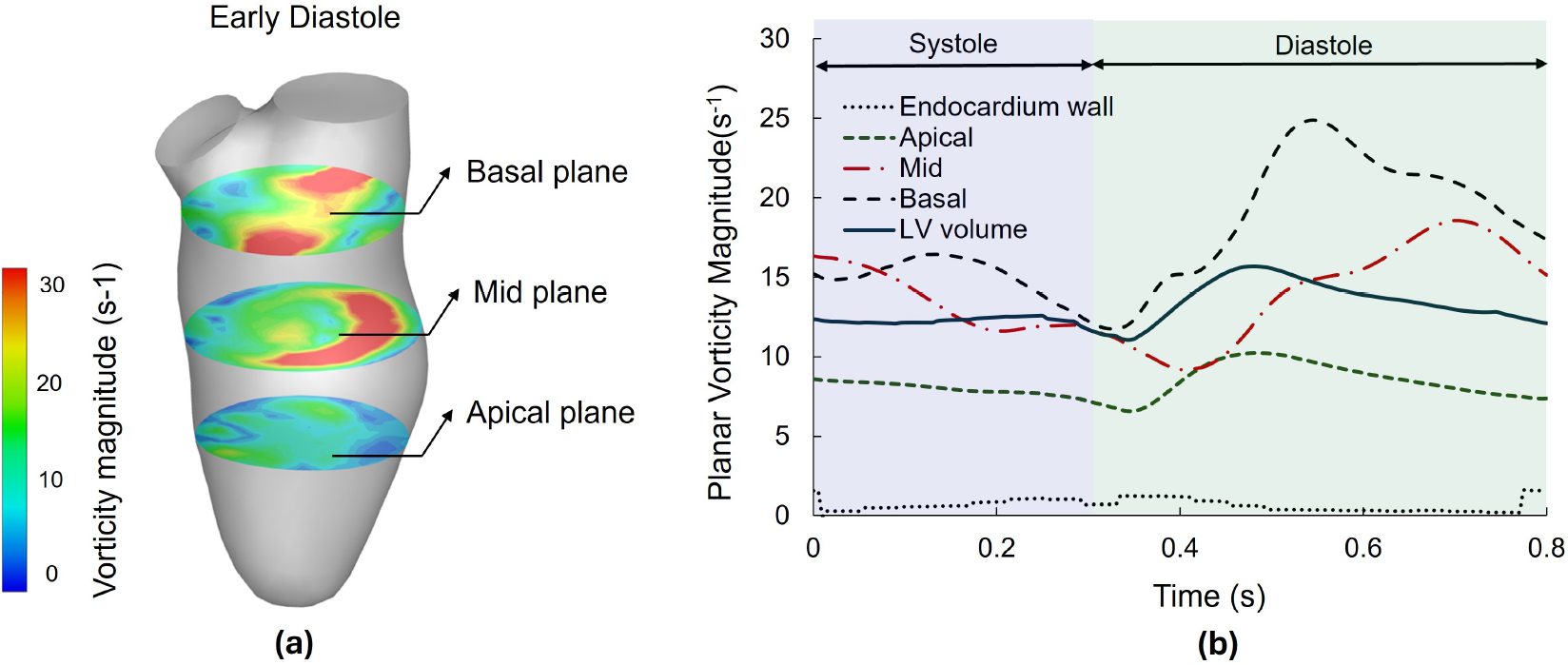
(a) Vorticity magnitude field at early systole phase in three short-axis planes of basal, mid, and apical sections. (b) Vorticity magnitude averaged in the three short-axis planes, the LV volume, and close to the endocardium wall during a cardiac cycle.

**Supplementary Figure S2.**
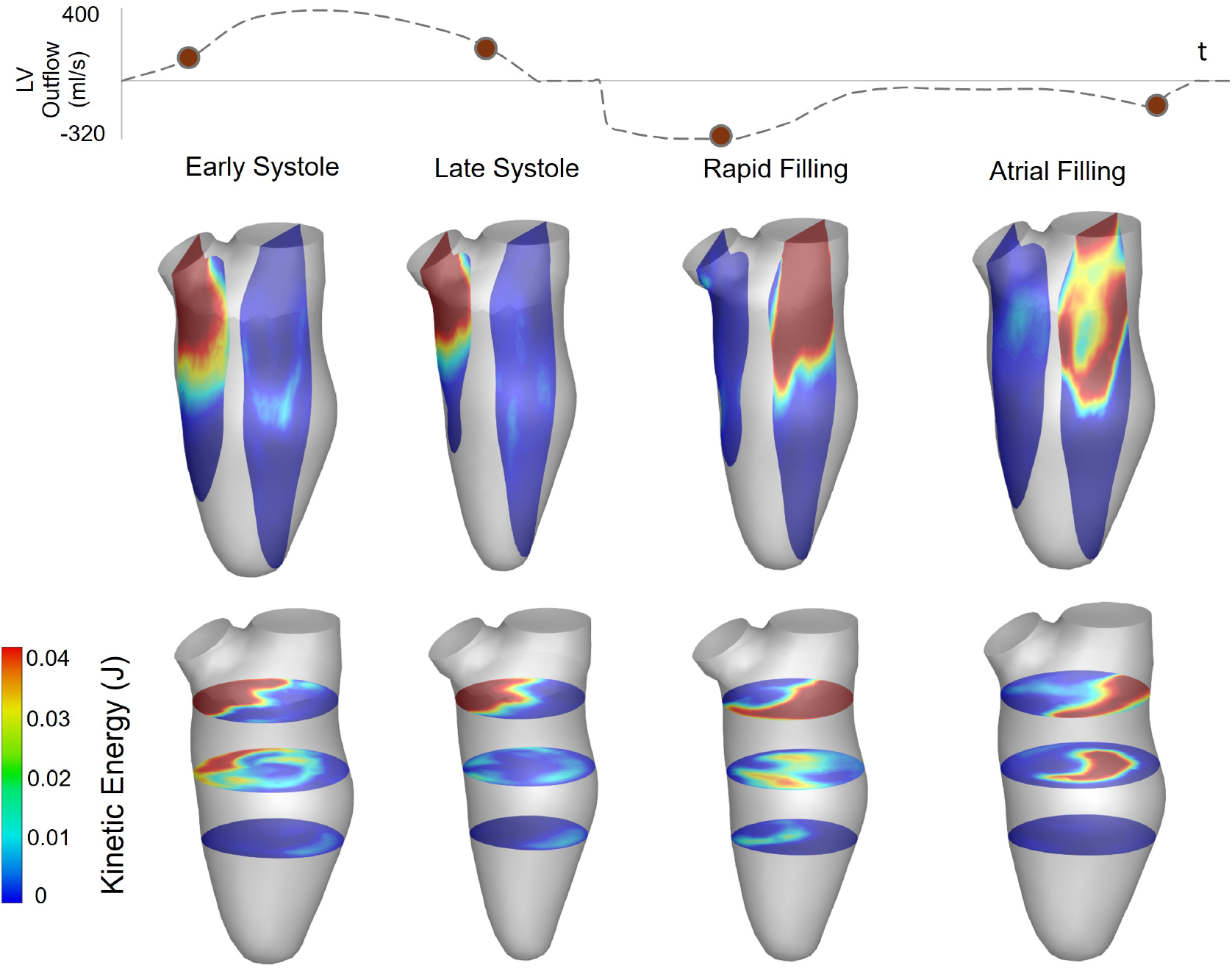
Kinetic energy field demonstrated at four time-points, early systole, late systole, rapid filling, and atrial filling, defined in the LV outflow vs time. The LVs at the top row illustrate kinetic energy at two parallel sagittal planes crossing the mitral and aortic orifices. The LVs at the bottom row illustrate the kinetic energy field at three SA planes, at basal, mid, and apical layers.

